# Decoding EEG for optimizing naturalistic memory

**DOI:** 10.1101/2023.08.25.553563

**Authors:** J.H. Rudoler, J.P. Bruska, W. Chang, M.R. Dougherty, B.S. Katerman, D.J. Halpern, N.B. Diamond, M.J. Kahana

## Abstract

**Background:** Spectral features of human electroencephalographic (EEG) recordings during learning predict subsequent recall variability.

**New method:** Capitalizing on these fluctuating neural features, we develop a non-invasive closed-loop (NICL) system for real-time optimization of human learning. Participants play a virtual navigation and memory game; recording multi-session data across days allowed us to build participant-specific classification models of recall success. In subsequent closed-loop sessions, our platform manipulated the timing of memory encoding, selectively presenting items during periods of predicted good or poor memory function based on EEG features decoded in real time.

**Results:** We observed greater memory modulation (difference between recall rates when presenting items during predicted good vs. poor learning periods) for participants with higher out-of-sample classification accuracy.

**Comparison with Existing Methods:** This study demonstrates greater-than-chance memory decoding from EEG recordings in a naturalistic virtual navigation task with greater real-world validity than basic word-list recall paradigms. Here we modulate memory by timing stimulus presentation based on noninvasive scalp EEG recordings, whereas prior closed-loop studies for memory improvement involved intracranial recordings and direct electrical stimulation. Other noninvasive studies have investigated the use of neurofeedback or remedial study for memory improvement.

**Conclusions:** These findings present a proof-of-concept for using non-invasive closed-loop technology to optimize human learning and memory through principled stimulus timing, but only in those participants for whom classifiers reliably predict out-of-sample memory function.

## 1 Introduction

Memory performance exhibits marked variability across time, as evident in momentary memory lapses that can cause us frustration or embarrassment. Sometimes, external factors trigger or interfere with our ability to learn and remember; for instance, the order in which items are presented affects the probability that they are subsequently recalled (Murdock, 1962; Deese & Kaufman, 1957). Some recall variability can also be attributed to item memorability or external contextual features (Aka, Phan, & Kahana, 2021; Rubin, 1985; Xie, Bainbridge, Inati, Baker, & Zaghloul, 2020). Recent work even shows that superior recall can be induced through model-based cue selection (Cornell, Norman, Griffiths, & Zhang, 2023). In addition to variability that can be predicted from exogenous variables, recent evidence suggests that at least some and possibly a large share of this variability arises endogenously (Kahana, Aggarwal, & Phan, 2018). One might expect that such endogenous variability is a function of observable neural activity – there is robust evidence that neural activity before and during memory encoding correlate with subsequent recall success, a phenomenon known as the subsequent memory effect (SME) (Sanquist et al., 1980; Paller & Wagner, 2002; Sederberg et al., 2003; Fell et al., 2011; Long et al., 2014; Griffiths et al., 2016; Rudoler et al., 2023). However, it is not yet clear if these patterns of neural activity that correlate with successful memory actually *cause* the observed variability (Weidemann & Kahana, 2021; Rubinstein, Weidemann, Sperling, & Kahana, 2023; Halpern, Tubridy, Davachi, & Gureckis, 2023).

The existence of the SME implies that we can reliably forecast successful memory encoding, and machine learning algorithms applied to both invasive and non-invasive recordings of brain activity have successfully predicted variation in memory performance at the level of both items (second-to-second) and lists (minute-to-minute) (Noh et al., 2014; Höhne et al., 2016; Astrand, 2018; Kragel et al., 2017; Phan et al., 2019; Chakravarty et al., 2020; Noh et al., 2018; Li et al., 2024; Weidemann et al., 2019; Weidemann & Kahana, 2021; Rubinstein et al., 2023). Accurately predicting downstream behavior based on neural activity laid the foundation for developing brain-computer interfaces for therapeutic intervention. In particular, recent studies have used model-based, closed-loop electrical stimulation of lateral temporal cortex to improve memory in neurosurgical patients (Ezzyat et al., 2018; Kahana et al., 2023).

Whereas these studies illustrate the promise of model-based closed-loop electrical stimulation as therapeutic intervention, other research has applied similar classification strategies to optimize learning without the use of electrical stimulation. One innovative study by deBettencourt et al. (2015) used real-time fMRI data to non-invasively train attention, though they did not directly examine the impact this might have on memory performance. In a follow-up study, they used fMRI-based neurofeedback to maximize contextual reinstatement. They showed that this procedure causally manipulated the probability of recalling items from a target reinstated context, as compared with items from an alternative context (deBettencourt, Turke-Browne, & Norman, 2019). Fukuda and Woodman (2015) derived electrophysiological biomarkers of encoding that predicted subsequent item recognition and found that “poorly-studied” items benefited from remedial study more than “well-studied” items. Another study unsuccessfully attempted to use a brain-computer interface to improve memory by optimizing the timing of item encoding (Burke, Merkow, Jacobs, Kahana, & Zaghloul, 2015). In that study, item presentation was triggered by univariate biomarkers associated with successful memory in earlier studies. Here, we build upon this approach by using regularized, multivariate regression models, which offer greater flexibility and predictive power. We also extend this approach to a more naturalistic recall paradigm.

The present study implemented a non-invasive closed-loop (NICL) procedure to optimize the timing of study-events to either improve or impair subsequent recall performance. Training multivariate classifiers on spectral EEG features, we manipulated the timing of study items to coincide with predicted good memory states (i.e., the set of brain states that predict encoding success) as opposed to bad memory states. We then examined whether these memory states causally affect subsequent memory performance. We hypothesized that classifier accuracy would modulate the efficacy of this procedure. Our study targeted a healthy population of young adults, and we embedded our memory task in a naturalistic virtual reality game with greater ecological validity than standard word-list tasks. Participants played the role of a bicycle courier, actively navigating through a virtual town to deliver objects to target locations (stores) and later attempting to recall all of the delivered objects. Dougherty et al. (in press) explore behavior in this task in detail and further discusses spatio-temporal memory dynamics.

We first designed a software platform for triggering item presentation based on classifier predictions from EEG features decoded in real time. Whereas prior work established the ability of classifiers to forecast out-of-session data, those studies retroactively used the full holdout session to normalize features, an approach incompatible with a real-time closed-loop system. Here, we demonstrate that classifiers are able to predict memory success in new sessions with just a modest amount of initial data used for normalization (though the success of this model generalization is highly variable across participants). Using models pre-trained on participant-specific task data, we then tested participants on separate days in closed-loop sessions where the timing of item encoding was triggered by predicted good encoding (‘optimize’ condition), predicted poor encoding (‘impair’ condition), or randomly (control condition).

Ultimately, we did not observe significant differences between conditions when averaging across the entire sample. However, since the closed-loop system is crucially dependent on the predictive model, we would expect the magnitude of differences between conditions to depend on the ability of the predictive model to classify mnemonic success. We therefore examined whether classifier performance moderated the success of the overall system, and found that performance correlates with recall differences across conditions in the expected direction. In other words, when we can reliably predict a given participant’s memory function, we can better manipulate it. These results point the way to building teaching systems that use the brain’s current state to optimize learning by presenting information to the user when it is most likely to stick.

## 2 Methods

### 2.1 Participants

A total of 49 participants consented to this study, but a number suffered from motion sickness, COVID exposures that interrupted the multi-day procedure, or technical issues with the closed loop technology (7 participants comprising the first round of testing), resulting in 21 participants who completed the entire experiment. All participants were young adults (ages 18–35) recruited among the students and staff at the University of Pennsylvania and neighboring institutions. The study was approved by the University of Pennsylvania Human Research Protections Program (HRPP), and all adult participants provided written informed consent to participate in this study.

### 2.2 Courier hybrid spatial-episodic memory task

Participants played the role of a courier in a hybrid spatial-episodic memory task, riding a bicycle and delivering parcels to stores located within a virtual town, as described above. Each experimental session consisted of a series of delivery days (i.e., trials), within which which participants navigated to a series of 15 stores. Participants were directed to each store with a text prompt (e.g. “Please find the grocery store”). Upon arriving at each store, after a 1±0.25 *s* jittered delay, they learned what objects they had delivered (via auditory and visual word presentation). In the standard version of the experiment, word presentation immediately followed this jittered delay. In sessions involving closed-loop intervention, the closed-loop system manipulated the timing and item presentation was dependent on classifier predictions (see *Closed-Loop Timining Manipulation*). At the end of each delivery day, participants were directed to a final 16th store, and upon arrival the screen went black and they were asked to freely recall that day’s delivered objects. During each recall phase in the experiment, we recorded vocal responses and subsequently annotated them offline using Penn Total Recall (https://memory.psych.upenn.edu/TotalRecall). For a detailed discussion of the methods underlying this memory task and analysis of spatio-temporal memory, we refer the reader to Dougherty et al. (in press).

**Figure 1.**
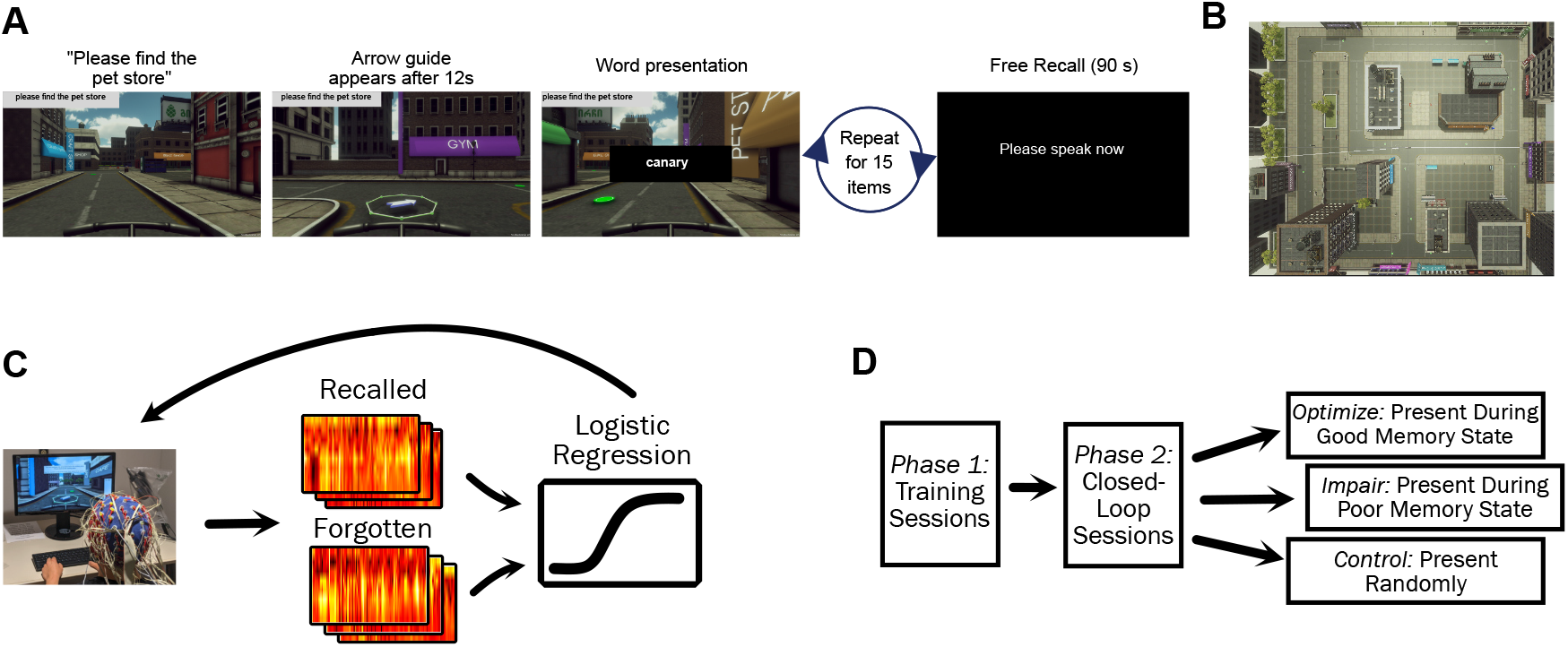
Paradigm. **A.** Participants performed a spatial navigation and episodic memory task in which they delivered items to various stores distributed about a virtual town. After a sequence of 15 deliveries, they had 90 seconds to recall as many of the delivered items as came to mind. **B.** Overhead view of the virtual town, which consisted of 17 stores and 18 non-store buildings. **C.** In a closed-loop paradigm, electrophysiological signals collected during the memory task were evaluated by a classifier in real-time to predict whether a studied item would be subsequently recalled or forgotten. These predictions explicitly affect gameplay, creating a feedback loop. **D.** Every participant completed 4-8 training sessions, followed by 2-4 closed-loop sessions. Within each closed-loop session, trials were split into 3 conditions in a randomized order: memory optimization (items presented when the participant was predicted to recall them), memory impairment (items presented when the participants was predicted to forget them), and a control (items presented randomly).

The experiment comprised two multi-session phases: During Phase 1, participants performed the hybrid spatial-episodic memory task - without any further manipulation - while we recorded 129-channel EEG signals. Phase 1 data was used to train participant-specific classifiers which discriminated between neural features predicting subsequent remembering and forgetting. In Phase 2, we evaluated closed-loop optimization of stimulus timing, to determine whether items studied during EEG-predicted “good-encoding states” resulted in better subsequent memory than items studied in EEG-predicted “poor-encoding states”. Specifically, we applied the classifiers to real-time neural activity, and modified the timing of item presentations based on classifier outputs (see *Closed-Loop Stimulus Presentation*). Each participant contributed between 5 and 8 Phase-1 recording sessions and 2-4 Phase-2 closed-loop sessions.

### 2.3 EEG recording and closed-loop infrastructure

We recorded electroencephalographic (EEG) data using a 128-channel BioSemi system with a 2048 Hz sampling rate. We used BioSemi’s proprietary software ActiView for data acquisition and monitoring the incoming data. BioSemi systems record data in the BioSemi Data Format (BDF), a 24-bit version of the popular European Data Format (EDF). Phase 1 sessions did not demand any use of the incoming data in real time, so we simply saved the data for later analysis. Importantly, these data were used to train classifiers that we subsequently employed during Phase 2 (closed-loop) sessions.

During Phase 2, we needed to modify the behavioral task in real time, based on changing neural features recorded in short, overlapping time intervals. To accomplish this, we split the incoming data stream so that we both saved a copy of the data to disk and also forwarded the data immediately to our own custom Python program for post-processing and analysis. The data pre-processing and feature extraction is described in the next section.

A primary challenge of implementing the closed-loop procedure is the need to analyze overlapping data epochs, since we are interested in changes of brain activity on a timescale which is shorter than the length of the data epoch (1 s) we analyze. This necessitates efficient parallelization of the data analysis for each epoch. As soon as the data for an epoch is collected, it is added to a queue which launches the analysis of that data in a parallel process as soon as resources are available. We thoroughly tested this system to ensure that the data collection rate did not in general exceed the speed of parallel analysis - if it did, the system would accumulate a backlog of data in the queue. If the data are not analyzed immediately, the classifier predictions become outdated. This system failure still occurred in a small subset of sessions that we excluded from our closed-loop memory improvement analyses (see Appendix A).

Once the spectral features of a data epoch were derived (a sub-200 ms process), we used the classifier trained on record-only data to predict the probability that the neural activity from that epoch represented a successful memory encoding state. This classifier result was then communicated from the EEG acquisition computer to the behavioral task computer by a TCP network connection. See Appendix A for detail about system performance and the typical duration of feature derivation and classifier prediction. EEG epochs of 1 s (the actual epoch is closer to 2 s, but we discard buffers to avoid edge artifacts) were collected every 250 ms – this means that we sample brain states at a rate of 4 Hz.

### 2.4 EEG Pre-processing and Spectral Decomposition

To generate training data for classifying the success of memory encoding in Phase 1, we constructed 1 second epochs spanning from the time window from 300 to 1300 ms following the onset of an item’s presentation. A prior closed-loop stimulation study found that training on brain activity during stimulus presentation led to effective closed-loop intervention with direct electrical stimulation (Ezzyat et al., 2018). The decision to train on EEG *during* stimulus presentation when we will ultimately classify brain states *prior* to presenting stimuli may seem unintuitive – however, the stimulus presentation period is when the neural features of successful encoding are strongest. The goal is to accurately identify these features in real-time and quickly present the next encoding stimulus. Training on EEG prior to stimulus onset is ineffective because the labels (recalled vs. not recalled) are heavily confounded by the neural features that arise while the stimulus is presented.

We applied a global average reference to the raw EEG recording, subtracting the average of 128 channels from the signal at each individual channel. A 1 Hz high-pass filter eliminated any residual channel-specific baseline drift in the signal, and a fourth-order Butterworth bandstop filter removed electrical line noise at 60 Hz as well as the 120 Hz harmonic. Next, we used the Morlet wavelet transform to estimate spectral power, convolving the EEG signal with Gaussian wavelets with a width corresponding to 5 cycles at the frequency of interest. We computed power separately for each channel at 8 logarithmically spaced frequencies from 6 to 180 Hz (6.00 Hz, 9.75 Hz, 15.86 Hz, 25.78 Hz, 41.90 Hz, 68.11 Hz, 110.73 Hz, 180.00 Hz). We included a buffer period on each side equal to half the wavelet width at the lowest frequency of interest (416.6 ms), in order to avoid contaminating the power estimate with edge effects, and discarded the buffer period from our time-resolved power estimates before proceeding. We excluded frequencies below 6 Hz from our analysis in order to preserve greater time resolution when later applying the classifier in real-time during Phase 2.

These power estimates yielded 1024 features - frequency-by-channel pairs representing every combination of 8 frequencies and 128 channels - for every time sample of every encoding event. We log-transformed these features and collapsed them by averaging over time to obtain a single value to represent each feature for every encoding event. We then *z*-scored the features - subtracting the mean and dividing by the standard deviation of all encoding events within each session - to normalize the data and standardize units across sessions. Normalization is an essential step for regularized regression because the scale of the regression weights heavily impacts the penalty term of the optimized loss function.

The EEG-processing was identical during Phase 2, with the exception of feature normalization. Since features had to be normalized in real time, instead of after data collection was completed, we could not *z*-score across all events within a still-in-progress session. Instead, we held out the first trial of each closed-loop session to establish a baseline for normalization.

### 2.5 Closed-Loop Stimulus Presentation

The term *closed-loop* refers to the feedback loop created by this experimental paradigm. A participant’s neural data collected during gameplay is used as input for a classifier that predicts the probability that a stimulus presented at the current time will be subsequently recalled. The classifier prediction is then used to manipulate gameplay (specifically, the timing of stimulus presentation for memory encoding) in the task, which indirectly influences that participant’s neural activity, which in turn is fed back into the classifier… and so on.

#### Classifier Training

Distinct participant-specific classifiers learned the neural features that best predicted memory for each person. We trained L2-penalized logistic regression classifiers based on item encoding events during record-only sessions: for each epoch corresponding to an encoded item, we regressed 1024 electrophysiological features (channel × frequency) against binary labels reflecting successful or failed retrieval during subsequent free recall. To ensure that the classes in the training data were balanced, we weighted the training instances in each class inversely proportional to the number of samples in the class. To select the L2 penalty parameter, we performed a leave-one-session-out cross validation and selected the penalty term with the best average accuracy score across held-out sessions. Then, we retrained a single classifier on the full training set using the selected penalty term and applied that classifier in closed-loop sessions.

#### Closed-Loop Timing Manipulation

During closed-loop sessions, our NICL system manipulated the delay prior to word presentation. This was achieved by pausing gameplay (i.e. during the delay between store arrival and item presentation; 1±0.25 *s* in record-only sessions) and presenting the awaited item contingent upon binary classification results (1 for predicted memory success and 0 for predicted memory failure). There were three different trial conditions which corresponded to different timing manipulations:

1. Memory optimization: wait until a prediction of 1 is received.
2. Memory impairment: wait until a prediction of 0 is received.
3. Control: wait a random amount of time sampled from a uniform distribution.

In addition to these three conditions, there was a trial at the beginning of each session without any timing manipulation that was used as a baseline for normalizing subsequently collected data. The timing of presentations on this trial was the same as in Phase 1: word presentation was delayed by 1±0.25 *s* after arriving at a store. When awaiting classifier results, the delay changed from the usual 1±0.25 *s* jitter to a 1 *s* delay and variable wait until the arrival of the desired classification result, with a timeout of 5 *s* (6 *s* total; the object was presented automatically at this point so the system did not wait for the target memory state indefinitely). Importantly, if the classifier presented at a “timeout”, the object is not actually presented in the target state. In the control condition, the delay changed from 1±0.25 *s* to a random delay sampled uniformly between 1 *s* and 6 *s*.

### 2.6 Statistical modeling of memory treatment effects

We are interested in modeling the causal effect that our closed-loop timing manipulation at encoding has on subsequent recall performance. By randomizing the timing condition (optimization, impairment, control) across lists, we control for confounders and are able to estimate the “treatment effect” directly (Greenland, 1990). As we predict the impairment, control, and optimization conditions to be ordered in terms of increasing recall, we encoded them as -1, 0, and 1 respectively. This allows us to test a single linear treatment effect instead of many pairwise comparisons between conditions. Given that the success of the timing manipulation hinges on the degree to which participants’ multivariate classifiers generalized from record-only to closed-loop sessions, we also investigated whether and how classifier accuracy shaped the NICL system’s influence on memory performance. To that end, we fit a linear mixed-effects model (Bates, Mächler, Bolker, & Walker, 2015) regressing average subsequent recall on trial type, closed-loop AUC, and their interaction (as in Figure 2B) with random intercepts for each participant. To ensure proper estimation of the effects and their standard errors, we initially tried to fit a maximal model that also included random slopes for both fixed effects and incrementally reduced the model complexity to remove zero-variance components and avoid singularities in the estimated variance-covariance matrix (Matuschek, Kliegl, Vasishth, Baayen, & Bates, 2017; Bates, Kliegl, Vasishth, & Baayen, 2018).

**Figure 2.**
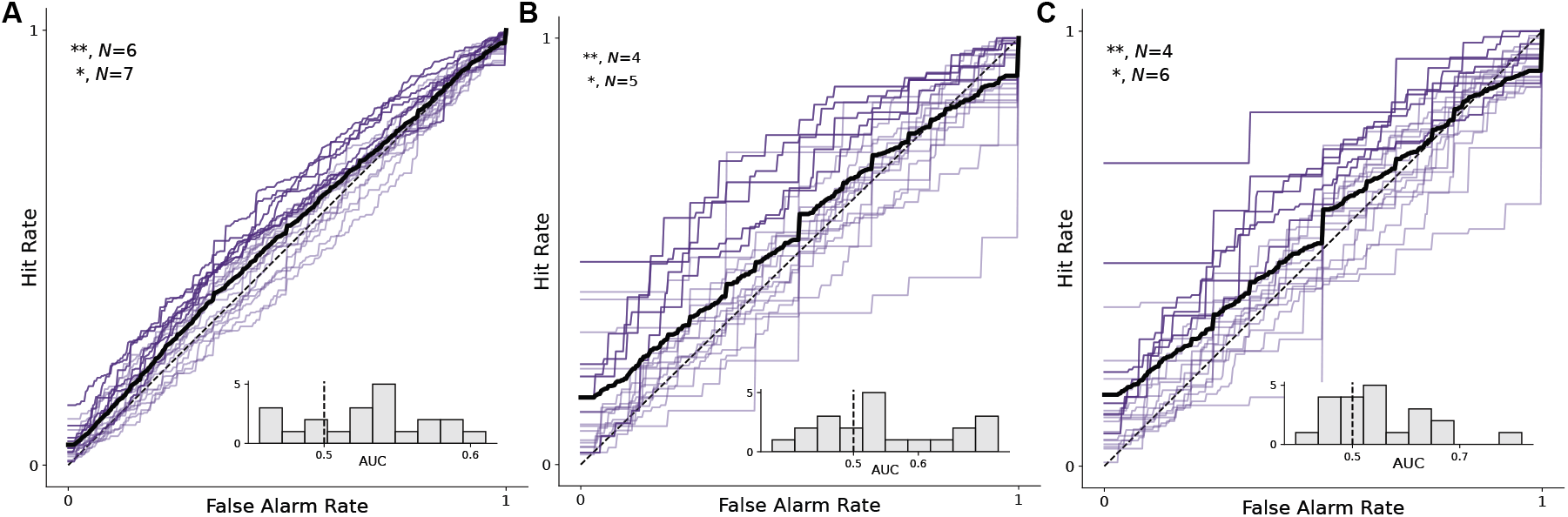
Predicting subsequent memory. Receiver Operating Characteristic (ROC) curves. Black line indicates average across *N* = 21 participants. Participants with above-chance prediction accuracy are marked with opaque lines while at-chance participants are transparent. For each participant, classifier significance was evaluated by comparing the observed AUC against a distribution of null AUCs obtained by randomly shuffling test labels (** gives the number of participants with *p < .*01, and * the number with *p < .*05). **A.** Performance on training data (using cross-validation to get and unbiased estimate). Classifiers are both trained and evaluated on neural features collected during stimulus presentation. **B.** In closed-loop sessions, again classifying successful encoding based on neural features stimulus item presentation. Performance in closed-loop sessions is evaluated only based on predictions for trials in which there is no systematic timing manipulation - normalization trials and control (random timing) trials. The conditions simulate real-time classification by using less data for normalization. **C.** Closed-loop performance, when classifying an epoch *prior* to stimulus onset (note that classifiers are still trained on data collected during stimulus presentation).

We report effects as *t*-values which represent the fixed effect estimate divided by the standard error of that estimate. The associated *p*-values arise from a *t*-test against the hypothesis that the effect size is equal to zero, using a corrected degrees of freedom term to account for possibly unequal sample sizes and variances (Satterthwaite, 1946; Kuznetsova, Brockhoff, & Christensen, 2017; Giesbrecht & Burns, 1985).

### 2.7 Data and Code Availability

Raw data and code for this study are freely available online. Behavioral and electrophysiological data comply with the Brain Imaging Data Standard (BIDS; Pernet et al., 2019; Appelhoff et al., 2019) and can be downloaded from OpenNeuro (Markiewicz et al., 2021) at https://openneuro.org/datasets/ds004706. Analysis code for all figures can be found at https://memory.psych.upenn.edu/Publications attached to the citation for this study, and additional scripts for data handling and preprocessing are publicly available at https://github.com/pennmem.

## 3 Results

A model-driven intervention depends on accurate prediction - we hypothesized, therefore, that our approach would improve memory *only if* the underlying classifiers could predict memory success above chance levels. In cases where classification is unreliable, we cannot evaluate the efficacy of our closed-loop optimization procedure. It is important to disambiguate failures to classify memory in the first place and failures to improve memory by optimizing encoding.

Our primary analyses focus on evaluating both our models and our closed-loop procedure. First, how well do machine learning classifiers trained on record-only sessions predict mnemonic success in closed-loop sessions? Second, how does the closed-loop task manipulation affect subsequent recall probability?

### Classifier Performance

We first trained participant-specific multivariate classifiers to distinguish encoding epochs of subsequently recalled and non-recalled items. Using a leave-one-session-out cross-validation scheme we computed the area under the Receiver Operating Characteristic function (“area under curve” or AUC) over the predicted recall probabilities of encoding events in the hold-out sessions. We evaluated the statistical robustness of an individual classifier via permutation testing: we randomly shuffled the test labels and recomputed the AUC 10,000 times, and set a threshold for above-chance classification at the 95th percentile of this distribution. This tests whether the observed performance could have been generated by chance, under the null assumption that the model has no useful information for predicting labels, since there is no meaningful statistical relationship between the features in the hold-out sets and their permuted labels. The data for both training and test in this analysis are derived from EEG collected during stimulus presentation (300-1300 ms after onset), as described in *EEG Pre-processing and Spectral Decomposition*. Figure 2A summarizes classifier performance across the training dataset and also shows data for each individual participant. At the group level (mean AUC = 0.529, SE = 0.01), the distribution of the observed AUCs significantly exceeded the chance-level score of 0.5 (*t*(21) = 3.09, *p* = 0.005).

Another desideratum is that recall scales with the probability of recall predicted by the classifiers - this provides a kind of sanity check that the predicted probabilities are meaningful. To this end we fit a linear mixed effects model, regressing the binary labels (recalled / not recalled) against the predicted probability for each item, with random slopes and intercepts for session nested within participant to account for individual differences. The relationship was indeed strong (*t*(21) = 2.57, *p* = 0.03) confirming that the actual recall rate scales with the classifier’s confidence.

Cross-validation in the training set, however, is only an estimate of classifier performance when generalizing to new sessions. Many sources of noise can change between sessions, causing a shift in the distribution of neural features for recalled and forgotten items. In closed-loop sessions, waiting for the target brain state extended the inter-stimulus intervals such that they ranged between 1 and 6 seconds – the performance of the classifiers may have been affected by this task change. Additionally, classifying in real-time precludes using the full session recording for data normalization. To evaluate the classifiers’ ability to generalize to the data distribution in the closed-loop sessions, it is necessary to test them in a way which accounts for these differences. To estimate real-time out-of-sample performance, we evaluated the predictions made on closed-loop data by the same classifiers which we used to conduct the closed-loop experiment. These classifiers were trained on data collected during stimulus presentation (300-1300 ms after onset) from all of the Phase 1 sessions for a given subject, using a penalty parameter chosen with cross-validation (see *Classifier Training* in *Methods*). As during training, they are also predicting recall based on EEG during stimulus presentation. In order to make sure that our results were unbiased by the classifier-driven manipulations, we evaluated predictions made on control trials with random timing of item presentation and on the trials which were held out as a baseline for normalization. We normalized the test data in the same manner as our closed-loop procedure – that is, by holding out the first list of each session and using it to compute a mean and standard deviation for *z*-scoring each feature (rather than computing the mean and standard deviation across the entire session). Figure 2B summarizes classifier performance on test data from closed-loop sessions (again, specifically for trials without the timing manipulation). At the group level (mean AUC = 0.549, SE = 0.02), the distribution of the observed AUCs significantly exceeded the chance-level score of 0.5 (*t*(21) = 2.26, *p* = 0.03). These performance scores are used for the subsequent analyses the following section, *Memory Improvement with Closed-Loop Stimulus Timing*.

There is one more potential problem for generalizing to closed-loop sessions: though classifiers learned the correlates of successful encoding by training on the stimulus presentation period, in closed-loop sessions they ultimately observed neural features *prior* to stimulus onset in order to make decisions about when to present the next stimulus. Accordingly, we also tried an even more conservative test of generalization – we repeated the same analysis, but instead of testing on the stimulus presentation period (300-1300 ms after onset) we chose a conservative window of 1616-616 ms before stimulus onset (to account for buffer time to collect and classify EEG) to demonstrate that the classifiers can predict encoding success even with such a delay. Figure 2C summarizes classifier performance on this data. At the group level (mean AUC = 0.548, SE = 0.02), the distribution of the observed AUCs significantly exceeded the chance-level score of 0.5 (*t*(21) = 2.30, *p* = 0.03).

### Memory Improvement with Closed-Loop Stimulus Timing

During Phase 2 (closed-loop) sessions we used classifiers trained during Phase 1 (record-only) sessions to control the timing of item presentation during memory encoding. Phase 2 sessions included three conditions, which varied across trials: control trials with random stimulus presentation, trials in which we *optimize* memory encoding by presenting stimuli when participants are in a predicted good memory state, and trials in which we *impair* memory encoding by presenting stimuli when participants are in a predicted poor memory state.

We predicted, a priori, that to the extent that classifiers reliably predicted successful memory, subjects would exhibit higher recall for ‘optimized’ timing lists than for ‘impaired’ timing lists, with the control condition falling between these two extrema. Without conditioning on classifier performance, our sample included subjects and sessions for whom the classifier failed to optimize the timing of stimulus presentation. Accordingly, Figure 3A shows that without controlling for this confound, average recall exhibits no appreciable differences by trial condition.

**Figure 3.**
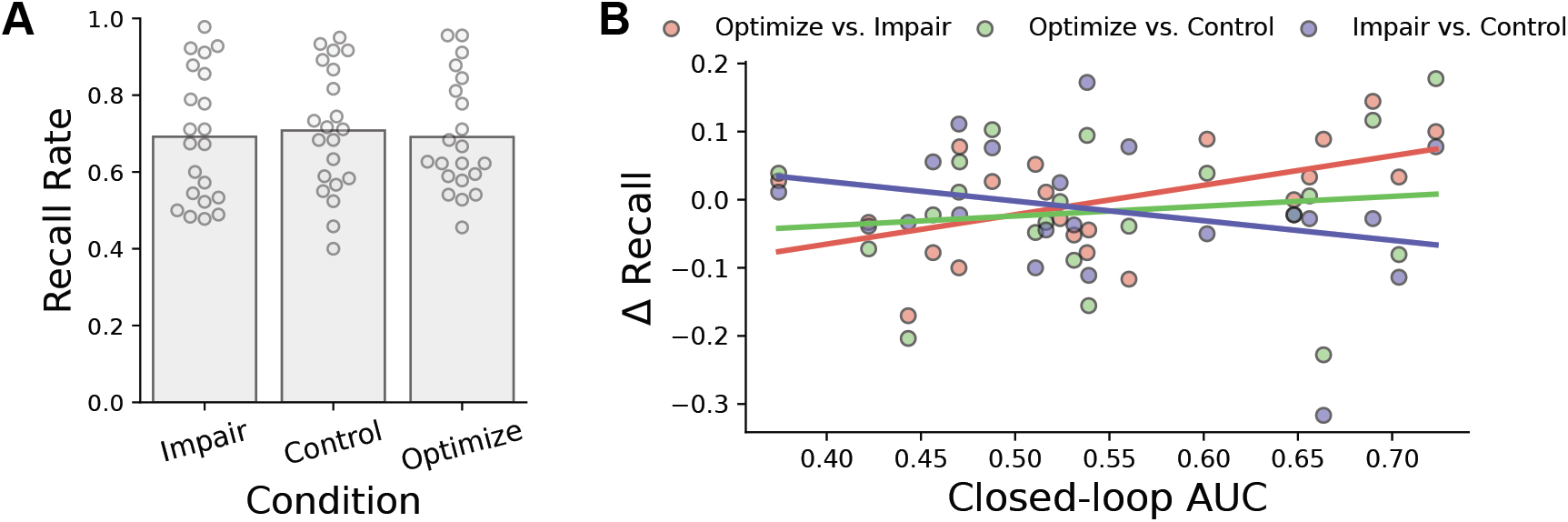
Recall and classifier performance. **A.** Average recall rate (the proportion of studied items that are subsequently recalled) for each trial condition in Phase 2. **B.** Pairwise differences in average recall rate as a function of classifier accuracy, with least-squares regression lines for each pair of conditions.

To test our predictions we fit a linear mixed-effects model to our data and estimated the effects of trial condition, classifier performance, and their interaction on average recall performance. This analysis did not reveal an overall main effect of trial condition on recall performance (i.e. the effect without regard to classifier accuracy). That is, there was no statistically significant difference in recall during trials with optimized timing across the sample of participants in our study, as compared with the control and the memory impairment conditions (see Figure 4A).

**Figure 4.**
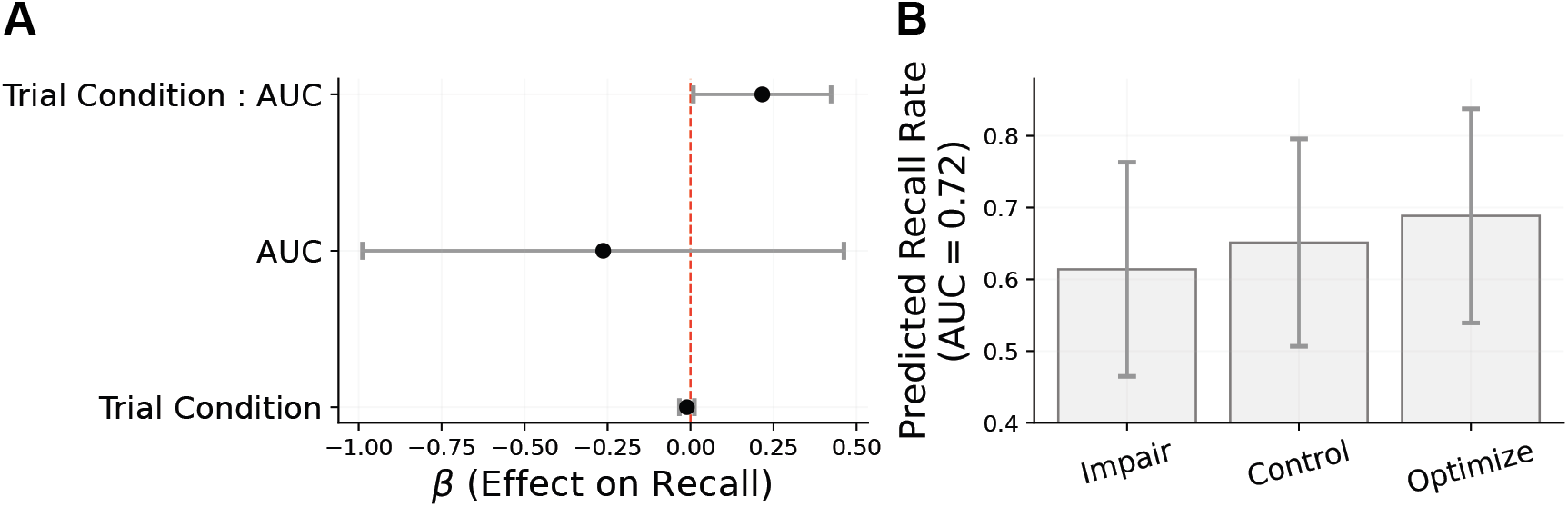
Modeling the effect of timing manipulations on subsequent recall. **A.** Linear mixed effects model coefficients for fixed effects, with 95% confidence intervals (testing against the null hypothesis that the effect size is zero). The model includes random intercepts for each participant, to account for individual differences in overall recall performance. **B.** Model-predicted recall rates for each trial condition at the highest observed classifier performance among participants in the study (AUC=0.72). Error bars represent 95% confidence intervals for the prediction, based on the standard errors of the estimates.

Consistent with our predictions, however, we find that treatment-induced differences in memory interact with participant-level classifier performance. Figure 3 shows pairwise differences in average recall by timing condition plotted against model performance on out-of-sample data. These relationships all trend in the expected directions – with greater predictive accuracy comes a larger memory boost on optimization trials as compared to impairment and control trials, along with a larger deficit for impairment trials as compared to control trials. Figure 4A shows the results from fitting a linear mixed effects model of subsequent recall with random intercepts for each participant. Model estimates show a significant positive correlation between average recall and the interaction of participant-level average AUC with trial condition (*t* = 2.11, *p* = 0.04). Figure 4B shows the model-predicted recall across trial conditions, at the highest observed out-of-sample AUC in our study. Though we fail to improve memory performance at the group level, we take this as an encouraging sign that a causal manipulation of subsequent memory may be possible with improved out-of-sample neural decoding.

As this is, to our knowledge, the first attempt to apply closed-loop scalp EEG-classifier methods to modulating human memory, we made various mis-steps along the way that should guide future work. Perhaps the most significant limitation evident from our analysis of the closed loop data was the generally poor level of classifier generalization between record-only and closed-loop sessions. The very nature of closed-loop experiments is that the classification is baked into the procedure, so even if one develops improved classification methods after the fact, evaluating these improvements requires collection of new experimental data. The next few sections evaluate various approaches we have taken to improving our classification methods. These approaches retroactively aid interpretation of the present data, and can guide future developments in scalp EEG-based closed-loop systems.

### Improving Classification

Closed-loop studies impose a high degree of scientific discipline on the researcher - a priori methodological choices by definition shape the unfolding of the closed-loop experiment, and therefore cannot be revised post hoc. Moreover, the technical constraints of the closed-loop architecture prevented us from applying certain standard methods used in scalp EEG to remove electromyographic artifacts, bad channels, etc. We thus sought to evaluate our pre-processing pipeline, to determine what changes might have led to superior out-of-sample classification performance. We compared cross-validated classifier performance on Phase 1 data with different pre-processing methods.

Our default pre-processing, as described in *Methods*, used a global average reference scheme coupled with a lowpass filter for drift and a bandpass filter to remove electrical line noise. In search of improvement, we first tried applying an ICA-driven approach called localized component filtering (DelPozo-Banos & Weidemann, 2017) to detect and remove artifacts caused by blinking or other noise sources. This approach decomposes EEG signals into their independent source components, applies artifact rejection algorithms locally (i.e. during specific time segments where artifacts appear), and then reconstructs artifact-free signals in EEG-sensor space. Using this “clean” EEG data instead of raw data did not improve classification (see Table D1, *t*(21)=0.193, *p*=0.85). We additionally tried applying a bipolar reference instead of a global average reference, in the hopes that a more finely resolved spatial filter would isolate informative features with specific spatial topography. Our choice of reference scheme did not have a significant impact on classifier accuracy either (see Table D1, *t*(21)=0.316, *p*=0.75).

A factor that we do know impacts classifier accuracy is data normalization. As discussed in *Methods*, we normalized our training features by applying a *z*-transform to each feature across all events within a session. This procedure, unfortunately, is impossible during the real-time closed-loop experiment: we do not yet have access to data from all events within the current session, as they have yet to be recorded. We negotiated a necessary trade-off between collecting sufficient normalization data and maximizing our valid experimental data: we first collect a subset of data (one trial or “delivery day”) from the current closed-loop session, and compute the mean and standard deviation of this subset in order to apply a real-time *z*-transform to the rest of the data as it is collected. This approach is unfortunately susceptible to low frequency drift and heteroscedasticity over the course of a recording session. As the local mean and variance of the features change, the *z*-transform will no longer correctly standardize the data to zero mean and unit variance. There is a fundamental trade-off between collecting more normalization data and collecting more closed-loop data, as the closed-loop procedure cannot begin until the normalization parameters have been set. Figure 5 shows how classifier accuracy reliably increases as we use more data for normalization (positive rate of increase is significant across participants; *t*(21)=3.19, *p*=.005). We further discuss the importance of normalization and look towards more advanced methods of aligning training and test domains in the *Discussion*.

**Figure 5.**
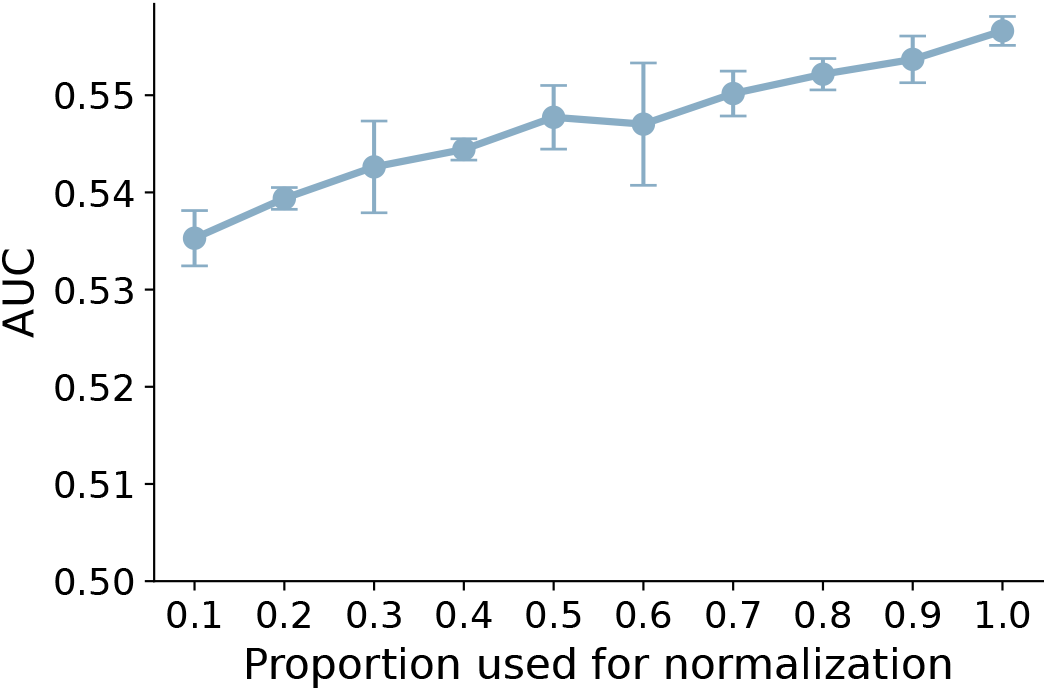
Feature normalization. Classifier performance in leave-one-session-out cross-validation on Phase 1 data improves as a function of the proportion of encoding events used in calculating the mean and standard deviation for the *z*-transform applied to the input features. Slope of increase is significant across participants (*t*(21)=3.19, *p*=.005). Error bars represent standard error of the mean across participants.

## 4 Discussion

We developed and evaluated a closed-loop platform that optimized the timing of memory encoding based on EEG classifiers trained to predict periods of good and poor memory function. We hypothesized that presenting memoranda during predicted good states - when the brain is ‘ready’ to encode new information - should lead to better memory than doing so during poor states. This hypothesis, however, hinges on the ability of trained classifiers to accurately forecast memory performance in closed-loop sessions. Evaluating this technology within the setting of an active virtual reality memory game afforded far greater ecological validity than prior studies using closed-loop brain-computer interface systems (Ezzyat & Suthana, in press; deBettencourt et al., 2015; Burke et al., 2015).

This proof-of-concept study led to three major conclusions supporting the viability of scalp EEG closed-loop timing to enhance human learning: First, we were able to design and implement a closed-loop system that applied multivariate decoding methods to EEG data in real-time, allowing our system to successfully control the timing of stimulus presentation based on decoded brain states. Second, we found that scalp EEG signals can reliably classify successful and unsuccessful learning on held-out sessions recorded on separate days, in a spatial-episodic memory task that provides far more complexity and real-world validity than previous experiments. Third, and most importantly, we found that the ability to decode memory states correlates with experimentally induced changes in recall performance.

We did not find reliable evidence of a population-level memory improvement - instead, we find that a successful demonstration of this type of closed-loop system’s overall causal effect on memory would require even more precision than achieved here. While this study obtained many more participants and sessions than earlier work, a crucial lesson for future work is that the resulting effects are likely to be too variable to be detected with the current design (see e.g. Figure 4b). One design flaw which we suspect impacted our results was choosing a presentation threshold for predicted recall probability (0.5) which treats the “good” and “bad” states as contiguous (see distributions in Appendix C); an alternative would be to identify “good” states above one threshold (e.g. 0.6) and “bad” states below another (e.g. 0.4) with a gap in between. The consequence of this choice is that many “good” and “bad” states are not so far apart in neural feature space, which might limit the behavioral differences caused by preferentially presenting in one or the other. It also means that the identified binary brain states change back and forth rapidly whenever the predicted probabilities are close to the threshold at 0.5. See in Appendix C that the stimuli were often presented almost immediately because these transitions between binary states happened so rapidly (or because the participant started off in the target state). As mentioned above in *Results*, we know that the continuous probability predictions do in fact scale with the actual recall rate – consequently we would expect that using a threshold to increase the difference in predicted probability between “good” and “bad” states should also correspond to greater recall differences.

Another major potential source of imprecision was poor classifier generalization, which, assuming that the mechanism for the potential success of such a closed-loop system was aligning stimulus timing to causally relevant brain states, should place limits on the possible effect sizes achievable. Specifically, although classifier performance was above chance in the aggregate, only a small proportion of the participants in our study exhibited sufficiently reliable generalization to closed-loop sessions.We see great potential in advancing these non-invasive closed-loop methods by improving our ability to build classifiers that generalize well to future recording sessions. Our data suggest that one key to improving classification is effective data alignment: bringing the training and test sets together into the same domain. Figure 5 shows that better normalization (via more data) leads to better classifier generalization.

A fundamental challenge of supervised machine learning is that the learning algorithm is trained on one dataset and subsequently asked to make predictions about an entirely non-overlapping dataset that can differ from the original in unpredictable or uncontrollable ways. Put another way, the classifier learns about one domain and is suddenly thrown into a different one: new noise sources, new signals, perhaps even new correlations between variables. Much can change between recording sessions: the precise locations of electrode contacts shift slightly, the impedances of the electrodes could differ, or an electrode might disconnect halfway through a session. A person’s underlying brain signals might change too: perhaps they had a sleepless night and forgot to have their morning coffee, or are distracted thinking about a final exam they have later in the week. Normalization is basically an attempt to put these potentially discrepant datasets on an equal footing. When normalized retroactively with already-collected full-session data, we demonstrate good classifier generalization to held out sessions recorded on different days. Thus, the challenge remains generalizing pre-trained classifiers to closed-loop data streaming into the classifier in real time.

This is an instance of a problem often referenced in the machine learning literature as *domain adaptation* – adapting a model trained in one situation to a new one. Domain adaptation is closely related to transfer learning, in which a pre-trained model is applied to a new task or problem. Our simplistic approach to this challenge was to apply a *z*-transform to the spectral features within each recording session; this transforms every feature onto a standard scale by subtracting the mean and dividing by the standard deviation. A limitation of this method is that it ignores correlations between features (i.e., it assumes that the data’s covariance matrix is diagonal). We know this not to be true - at the very least there are spatial correlations among neighboring electrode contacts that pick up on the same brain signals. Some simple domain adaptation algorithms (e.g. Sun, Feng, & Saenko, 2015) exploit the full covariance structure of the data and thereby account for more of the differences between learned domains. *z*-scoring across chronologically ordered events is also assumes the data are stationary, another assumption that is in all likelihood violated in any EEG data. Future work should explore time-sensitive, or adaptive, normalization techniques. Another promising area for future work is to combine training data across participants and thereby assemble a much larger training set. This would prevent overfitting an individual participant’s data and likely improve classification accuracy. Recent work demonstrates that using self-supervised representation learning to pre-train neural decoding models on large, heterogenous datasets may enable robust and efficient generalization to new domains (Azabou et al., 2023; Wang et al., 2023).

Of theoretical importance – if our NICL system had successfully manipulated memory encoding success, these findings would provide novel causal evidence for the importance of endogenous neural fluctuations in shaping memory success (Weidemann & Kahana, 2021; Halpern et al., 2023; Rubinstein et al., 2023). Decades of studies on the subsequent memory effect (SME) show how brain states at the time of item presentation differ between successful vs. unsuccessful memory encoding. We inverted this design here, which allowed for testing whether the fate of a discrete learning experience (i.e. a given stimulus presentation) is caused by the ‘brain time’ at which it is presented. The correlation between our ability to decode such states and the differences in recall performance hint at such a mechanism underlying the results.

While this study did not produce the targeted group-level memory improvement, careful analysis points the way forward to improved closed-loop systems and studies of those systems. As machine learning methods continue to advance quickly, along with increasingly affordable high-performance computing that can alleviate some computational challenges discussed in the appendix, we are optimistic that future work will realize the potential for memory improvement with brain-computer interfaces.

## Acknowledgements

### Contributions

J.H.R., M.J.K., D.J.H, and N.B.D. contributed to the final manuscript. J.H.R. and D.J.H. contributed to data analysis. J.H.R. and J.P.B. contributed to engineering the closed-loop recording and feedback system. M.J.K. and N.B.D. contributed to study conception and design. W.C., M.D.R., and B.S.K. contributed to data collection and study logistics.

### Funding

The authors gratefully acknowledge support from the U.S. Army Medical Research and Development Command (USAMRDC) through the Medical Technology Enterprise Consortium (MTEC) project MTEC-20-06-MOM-013, “Restoring memory with task-independent semi-chronic closed-loop direct brain stimulation and non-invasive closed-loop stimulus timing optimization”.

### Correspondence

Any questions or other correspondence concerning this manuscript should be addressed to Michael J. Kahana.

## Appendix A Evaluating closed-loop system performance

Imperative to the success of the NICLS experimental paradigm is the efficiency of computing and communicating real-time classifier predictions. We found that system performance was generally good, with most classifier results computed within about 200 ms. In some cases, though, the system failed unpredictably and computation lagged, leading to a compounding backlog of unprocessed data. When this happens, the classifier predictions are not communicated back to the behavioral task in time to be actionable.

**Figure A1.**
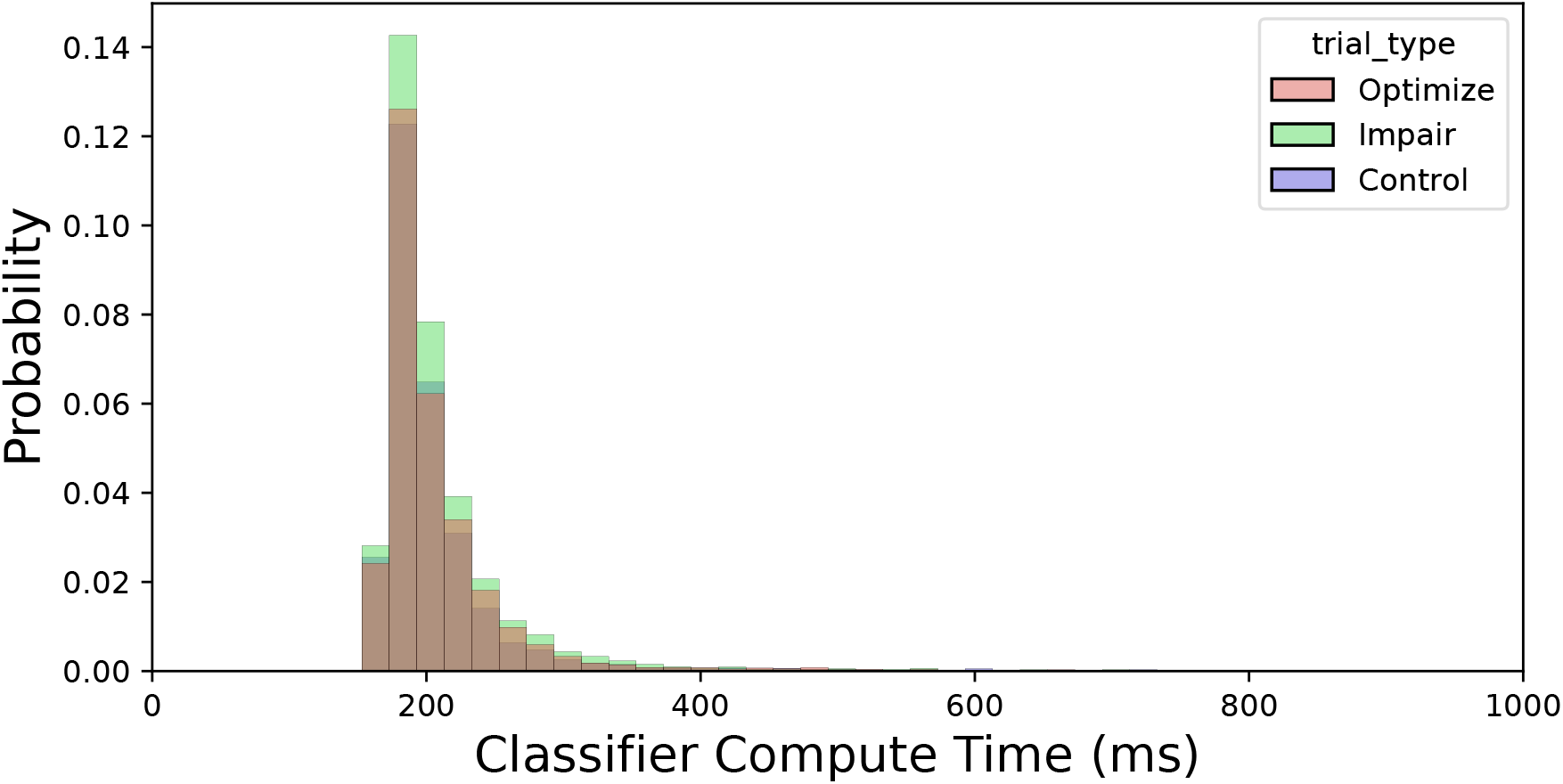
Distribution of durations for classifier computation.

Table A1 catalogs system performance for each participant and session in our closed-loop dataset. The maximum duration of time after an EEG epoch is collected that a classifier result can still be used to manipulate the task - before it “times out” and the item is presented anyway – is 6 seconds. We labeled any classifier results that took this long to compute as “overtime” and marked any sessions where this happened at least 5% of the time. These sessions were excluded from our data analysis (7 sessions total).

**Table A1.**
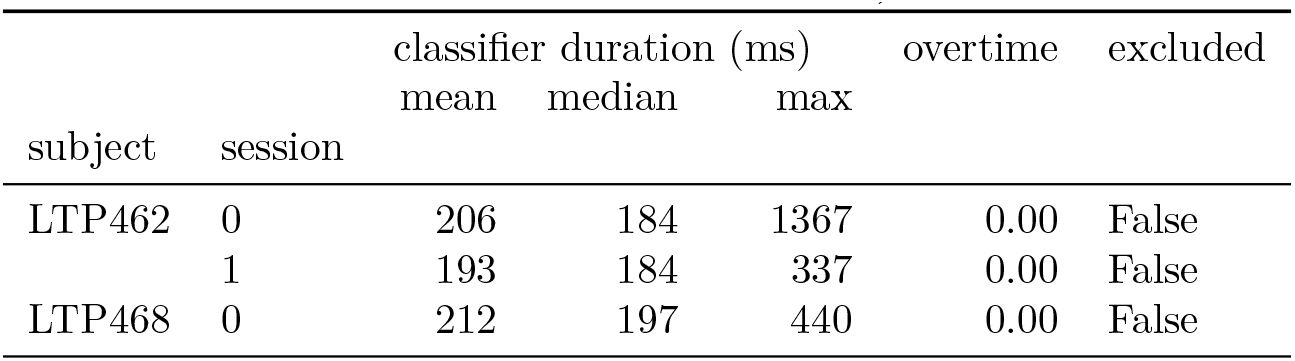
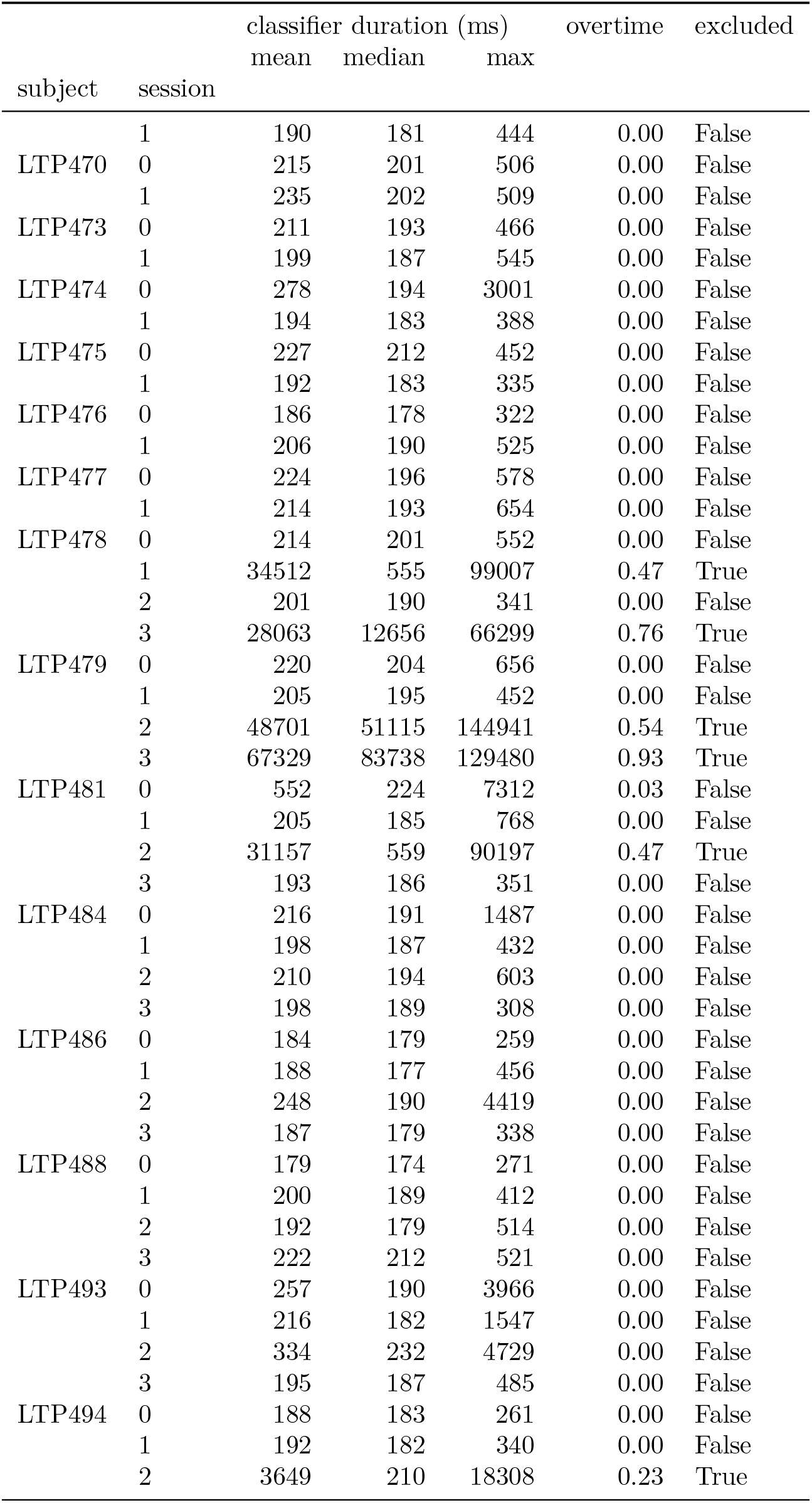
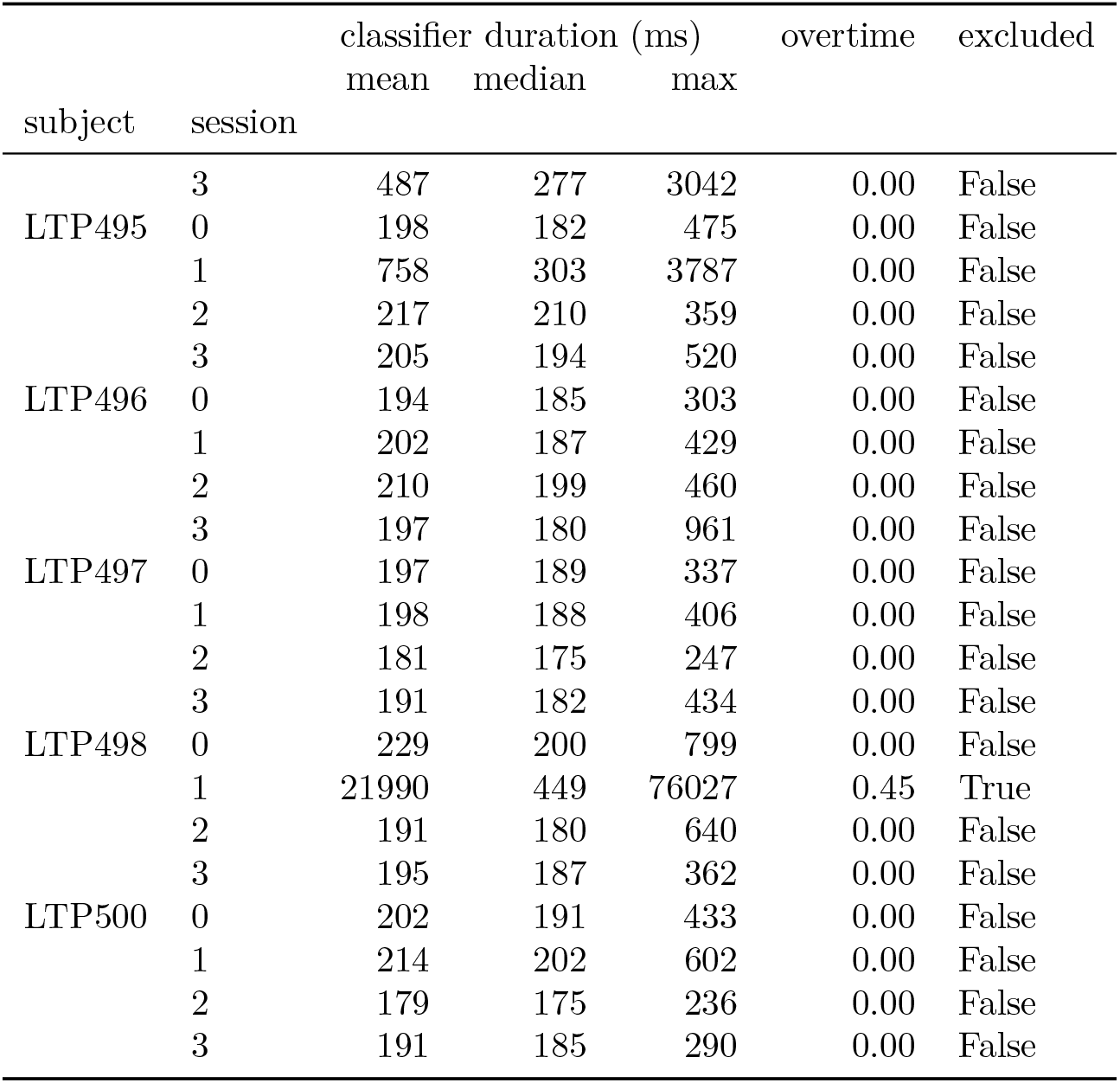
System Performance. Data for each subject and session in the closed-loop dataset, detailing how long computation of classifier predictions lasted and whether or not the duration of that computation exceeded our tolerance threshold of 6 seconds (the maximum compute time that would still allow the prediction to be actionable)

## Appendix B Timescale of classifier transitions

Classifier predictions, like the neural activity which the classifiers receive as input, are highly autocorrelated. After all, neural features are continuous: we have artificially binarized them. So, it takes some nonzero amount of time to transition from neural features associated with failed encoding to one associated with successful encoding. One might ask approximately how long it takes for “memory states” to transition. Figure B1 shows the distribution of time that the NICLS task spent waiting for the arrival of the target classifier results. Note the heavy weight close to zero - this reflects that at the time the encoding period began, the participant was already in the target memory state. Note that if a given participant’s classifier predicted a good/bad memory encoding state half the time, on average, then items would be presented immediately roughly half the time.

**Figure B1.**
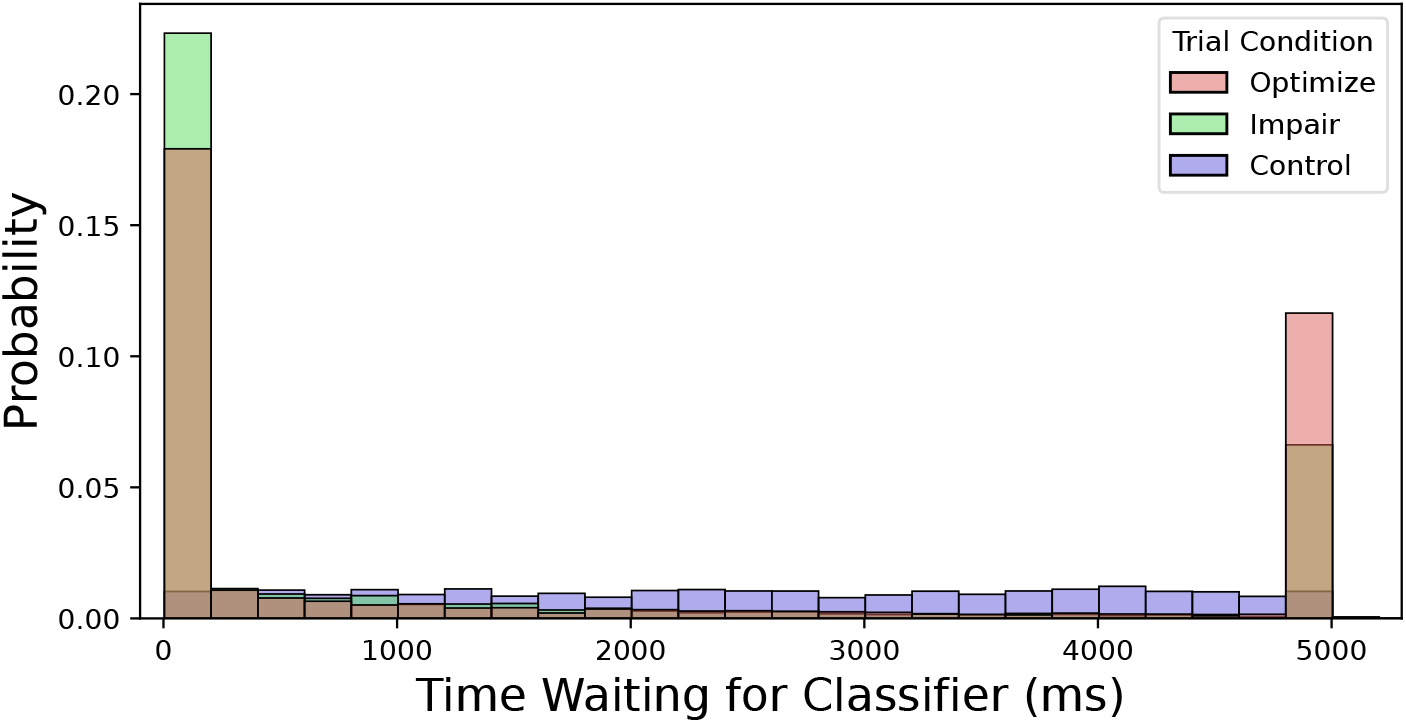
Distribution of time spent waiting for desired classifier result, split by trial condition.

## Appendix C Distribution of classifier predictions

We expect the distribution of classifier predictions that triggered a presentation to differ by trial condition, since they are explicitly constrained: “Optimize” trials only trigger presentation for predictions greater than 0.5 and “Impair” trials only trigger presentation for predictions below 0.5. “Control” trials are unconstrained; they present randomly, so the associated predictions should represent the true underlying distribution of model predictions.

Notice in Figure C1, predictions are densely centered near 0.5 – meaning, the model cannot distinguish class membership with confidence. This low confidence is an unintended consequence of choosing the experiment’s decision threshold for item presentation to be 0.5. That is to say, because neural activity is autocorrelated, if the experiment starts waiting in the non-target state (e.g. waiting for a good memory state [>0.5] and the participant is currently in a bad memory state [<0.5]) then the first prediction in the target state will likely be close to the threshold.

There are also peaks at 0 and 1, representing predictions made with extremely high confidence. This could be good or bad: for example, this could happen if the electrode cap moves too much and distribution of neural features shifts dramatically from the learned decision boundary, so that many or all events are predicted to belong to one class with high confidence.

**Figure C1.**
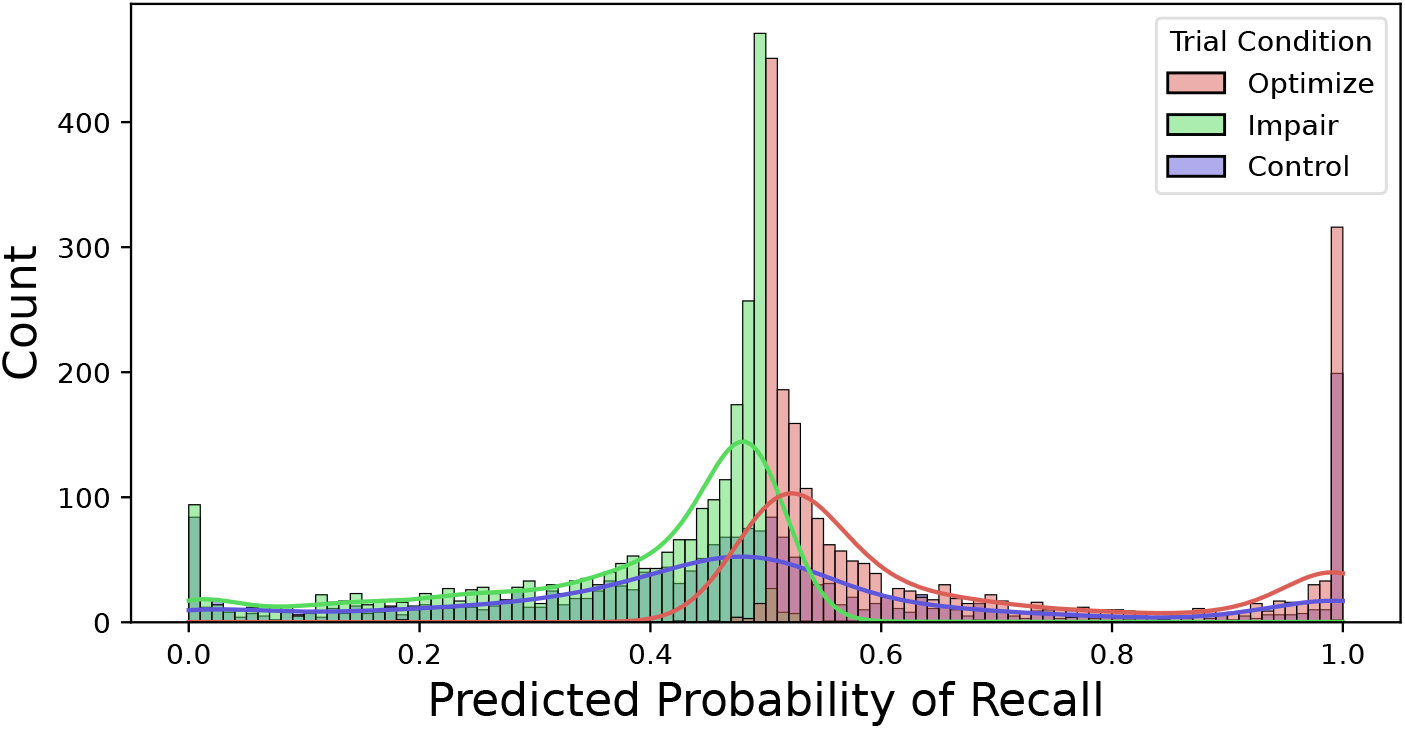
Distribution of classifier prediction values which triggered an item presentation, split by trial condition.

## Appendix D

**Table D1.**
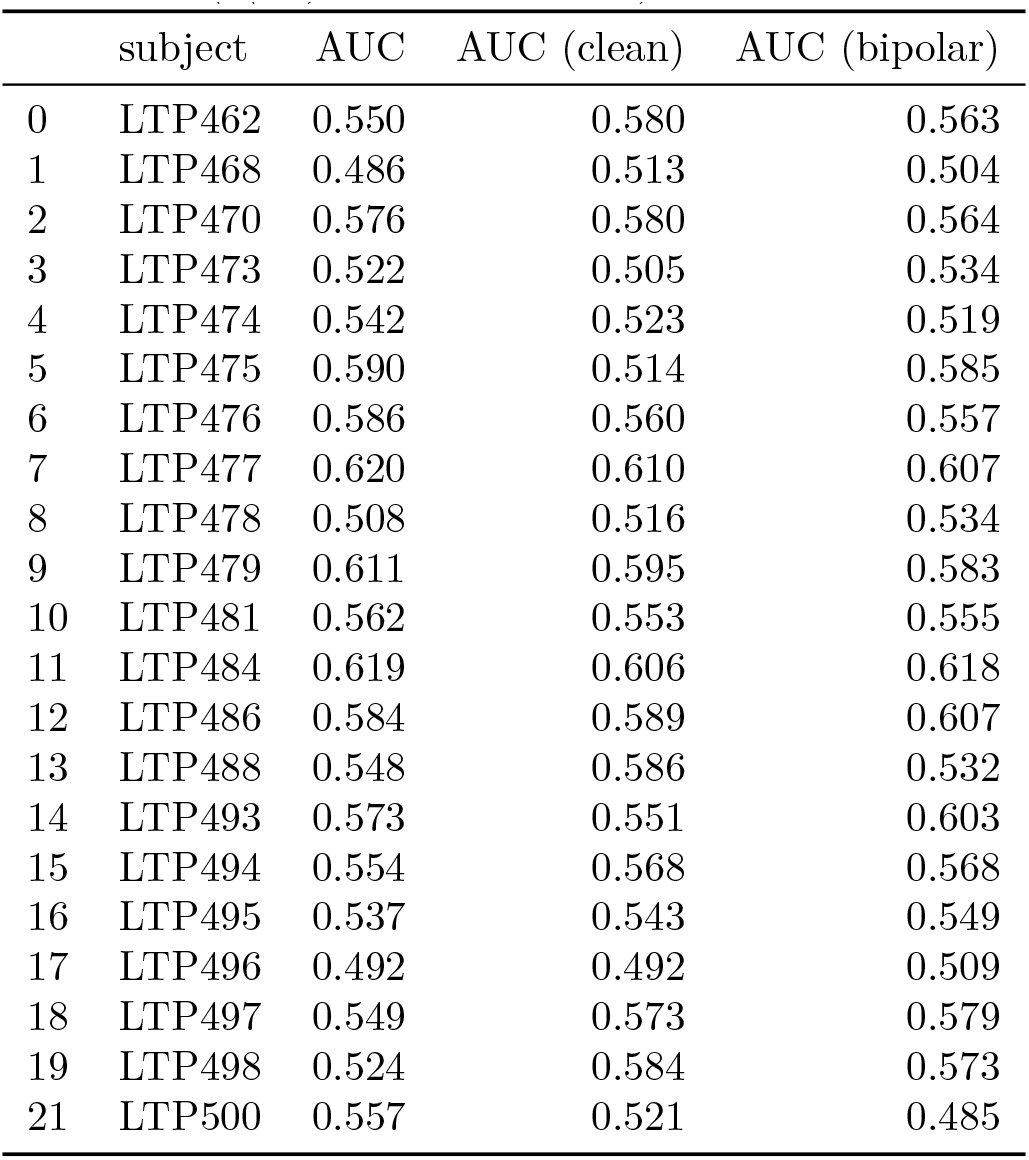
Preprocessing Methods and Classifier Performance. We found no significant difference in classifier performance (AUC) between different pre-processing methods. Our default pre-processing was to just use a global average reference scheme coupled with a lowpass filter for drift and a bandpass filter to remove electrical line noise. We compared this approach to pre-processing with ICA-based localized component filtering (t(21)=0.193, p=0.85) and also to a bipolar reference scheme (t(21)=0.316, p=0.75).

## Appendix E Non-Invasive Closed Loop (NICL) System Design and Flaws

### 6.1 Closed Loop Design

The closed-loop design is split into two separate programs: the NiclServer and the Task. The **NiclServer** is a backend which handles data flow from the BioSemi equipment, preprocesses the EEG, and computes a classifier prediction. The **Task** is the virtual reality game described above, which only differs from the original Courier game in that it listens for TCP messages from the NiclServer and updates an internal boolean with the most recent classifier result. Each components of NICLS is run on a separate computer, as both are computationally expensive. Our lab’s standard scalp data collection setup already uses two computers, to alleviate the computational burden of both running a virtual reality task and collecting high-density neural recordings.

The closed-loop system is split into two phases: normalization and classification.

#### Normalization

The system starts in the normalization phase, during which it computes a mean and standard deviation for each spectral feature for a series of encoding events. This mean and standard deviation are used to *z*-score subsequent encoding events before classifying it. The task waits until each encoding period and then sends a network message to NiclServer that an encoding took place. NiclServer receives the message and then updates a rolling mean and standard deviation with the corresponding EEG epoch.

#### Classification

In NiclServer, a classification happens every 125ms using the last 2 seconds of EEG data. This classification then sends a network message to the Task indicating the boolean state of classification. The task uses a separate thread to maintain a boolean of the most recent state of classification. It then uses the boolean for the closed-loop stimulus presentation during the task’s encoding and cued retrieval phase.

### 6.2 Limitations

While it does not affect the majority of our experimental data, we include this cautionary note as a warning for future system designers. The system uses two threads in a process pool in order to run the classifications in parallel. This is effective for speeding up the number of classifications per second, but the implementation does not guarantee that the classifications would be finished and reported in the correct order. This is not very problematic in our case for two reasons: there were only two parallel processes (causing a 125ms error) and the task decisions only relied on one value to be good or bad. If there were more processes running at the same time (with the same number of classifications per second) or if a task action required a number of the same classification decisions in a row, the potential for errors would greatly increase.

## References

Aka, A., Phan, T. D., & Kahana, M. J. (2021). Predicting recall of words and lists. Journal of Experimental Psychology: Learning, Memory, and Cognition, 47 (5), 765–784. doi: 10.1037/xlm0000964

Appelhoff, S., Sanderson, M., Brooks, T. L., van Vliet, M., Quentin, R., Holdgraf, C., … Jas, M. (2019). Mne-bids: Organizing electrophysiological data into the bids format and facilitating their analysis. Journal of Open Source Software, 4 (44), 1896. Retrieved from 10.21105/joss.01896 doi: 10.21105/joss.01896

Astrand, E. (2018, April). A continuous time-resolved measure decoded from eeg oscillatory activity predicts working memory task performance. Journal of Neural Engineering, 15 (3).

Azabou, M., Arora, V., Ganesh, V., Mao, X., Nachimuthu, S., Mendelson, M., … Dyer, E. (2023). A unified, scalable framework for neural population decoding. In Neurips.

Bates, D., Kliegl, R., Vasishth, S., & Baayen, H. (2018). Parsimonious mixed models. arXiv(arXiv:1506.04967). doi: 10.48550/arXiv.1506.04967

Bates, D., Mächler, M., Bolker, B., & Walker, S. (2015). Fitting linear mixed-effects models using lme4. Journal of Statistical Software, 67 (1), 1–48. doi: 10.18637/jss.v067.i01

Burke, J. F., Merkow, M., Jacobs, J., Kahana, M. J., & Zaghloul, K. (2015). Brain computer interface to enhance episodic memory in human participants. Frontiers in Human Neuroscience, 8, 1055.

Chakravarty, S., Chen, Y. Y., & Caplan, J. B. (2020). Predicting memory from study-related brain activity. Journal of Neurophysiology, 124, 2060–2075.

Cornell, C. A., Norman, K. A., Griffiths, T. L., & Zhang, Q. (2023). Improving memory search through model-based cue selection. PsyArXiv.

deBettencourt, M. T., Cohen, J. D., Lee, R. F., Norman, K. A., & Turk-Browne, N. B. (2015). Closed-loop training of attention with real-time brain imaging. Nature Neuroscience, 18 (3), 470–475.

deBettencourt, M. T., Turke-Browne, N. B., & Norman, K. A. (2019, October). Neuro-feedback helps to reveal a relationship between context reinstatement and memory retrieval. NeuroImage, 200, 292–301.

Deese, J., & Kaufman, R. A. (1957). Serial effects in recall of unorganized and sequentially organized verbal material. Journal of Experimental Psychology, 54, 180–187. doi: 10.1037/h0040536

DelPozo-Banos, M., & Weidemann, C. T. (2017). Localized component filtering for electroencephalogram artifact rejection. Psychophysiology, 54 (4), 608–619.

Dougherty, M. R., Chang, W., Rudoler, J. H., Katerman, B. S., Halpern, D. J., Bruska, J. P., … Kahana, M. J. (in press). Neural correlates of memory in a naturalistic spatiotemporal context. Journal of Experimental Psychology: Learning, Memory, and Cognition. doi: 10.1101/2022.11.30.518606

Ezzyat, Y., & Suthana, N. (in press). Oxford handbook of human memory. In M. J. Kahana & A. D. Wagner (Eds.), (2nd ed., chap. Brain Stimulation). Oxford, U. K.: Oxford University Press.

Ezzyat, Y., Wanda, P., Levy, D. F., Kadel, A., Aka, A., Pedisich, I., … Kahana, M. J. (2018). Closed-loop stimulation of temporal cortex rescues functional networks and improves memory. Nature Communications, 9 (1), 365. doi: 10.1038/s41467-017-02753-0

Fell, J., Ludowig, E., Staresina, B., Wagner, T., Kranz, T., Elger, C. E., & Axmacher, N. (2011). Medial temporal theta/alpha power enhancement precedes successful memory encoding: evidence based on intracranial eeg. Journal of Neuroscience, 31 (14), 5392–5397.

Fukuda, K., & Woodman, G. F. (2015). Predicting and improving recognition memory using multiple electrophysiological signals in real time. Psychological science, 26 (7), 1026–1037.

Giesbrecht, F. G., & Burns, J. C. (1985, June). Two-stage analysis based on a mixed model: Large-sample asymptotic theory and small-sample simulation results. Biometrics, 41 (2), 477. Retrieved from 10.2307/2530872 doi: 10.2307/2530872

Greenland, S. (1990). Randomization, statistics, and causal inference. Epidemiology, 1 (6).

Griffiths, B., Mazaheri, A., Debener, S., & Hanslmayr, S. (2016). Brain oscillations track the formation of episodic memories in the real world. NeuroImage, 143, 256–266.

Halpern, D. J., Tubridy, S., Davachi, L., & Gureckis, T. M. (2023). Identifying causal subsequent memory effects. Proceedings of the National Academy of Sciences.

Höhne, M., Jahanbekam, A., Bauckhage, C., Axmacher, N., & Fell, J. (2016, October). Prediction of successful memory encoding based on single-trial rhinal and hippocampal phase information. NeuroImage, 139, 127–135.

Kahana, M. J., Aggarwal, E. V., & Phan, T. D. (2018). The variability puzzle in human memory. *Journal of Experimental Psychology: Learning*, Memory, and Cognition, 44 (12), 1857–1863. doi: 10.1037/xlm0000553

Kahana, M. J., Ezzyat, Y., Wanda, P. A., Solomon, E. A., Adamovich-Zeitlin, R., Lega, B. C., … Diaz-Arrastia, R. R. (2023, July). Biomarker-guided neuromodulation aids memory in traumatic brain injury. Brain Stimulation, 16 (4), 1086–1093. doi: 10.1016/j.brs.2023.07.002

Kragel, J. E., Ezzyat, Y., Sperling, M. R., Gorniak, R., Worrell, G. A., Berry, B. M., … Kahana, M. J. (2017). Similar patterns of neural activity predict memory function during encoding and retrieval. NeuroImage, 155, 60–71. doi: 10.1016/j.neuroimage.2017.03.042

Kuznetsova, A., Brockhoff, P. B., & Christensen, R. H. B. (2017). lmerTest package: tests in linear mixed effects models. Journal of Statistical Software, 82 (13), 1–26. doi: 10.18637/jss.v082.i13

Li, Y., Pazdera, J. K., & Kahana, M. J. (2024). Eeg decoders track memory dynamics. Nature Communications, 15 (1), 2981. doi: 10.1038/s41467-024-46926-0

Long, N. M., Burke, J. F., & Kahana, M. J. (2014). Subsequent memory effect in intracranial and scalp EEG. NeuroImage, 84, 488–494. doi: 10.1016/j.neuroimage.2013.08.052

Markiewicz, C. J., Gorgolewski, K. J., Feingold, F., Blair, R., Halchenko, Y. O., Miller, E., … Poldrack, R. (2021, oct). The openneuro resource for sharing of neuroscience data. eLife, 10, e71774. Retrieved from 10.7554/eLife.71774 doi: 10.7554/eLife.71774

Matuschek, H., Kliegl, R., Vasishth, S., Baayen, H., & Bates, D. (2017). Balancing type i error and power in linear mixed models. Journal of Memory and Language, 94, 305–315. doi: 10.1016/j.jml.2017.01.001

Murdock, B. B. (1962). The serial position effect of free recall. Journal of Experimental Psychology, 64 (5), 482–488. doi: 10.1037/h0045106

Noh, E., Herzmann, G., Curran, T., & de Sa, V. R. (2014). Using single-trial eeg to predict and analyze subsequent memory. NeuroImage, 84, 712–723.

Noh, E., Liao, K., Mollison, M. V., Curran, T., & de Sa, V. R. (2018). Single-trial eeg analysis predicts memory retrieval and reveals source-dependent differences. Frontiers in Human Neuroscience, 12, 258.

Paller, K. A., & Wagner, A. D. (2002). Observing the transformation of experience into memory. Trends in Cognitive Sciences, 6 (2), 93–102. doi: 10.1016/S1364-6613(00)01845-3

Pernet, C. R., Appelhoff, S., Gorgolewski, K. J., Flandin, G., Phillips, C., Delorme, A., & Oostenveld, R. (2019). Eeg-bids, an extension to the brain imaging data structure for electroencephalography. Scientific Data, 6 (1), 103. Retrieved from 10.1038/s41597-019-0104-8 doi: 10.1038/s41597-019-0104-8

Phan, T. D., Wachter, J. A., Solomon, E., & Kahana, M. J. (2019). Multivariate stochastic volatility modeling of neural data. eLife, 8, e42950.

Rubin, D. C. (1985). Memorability as a measure of processing: a unit analysis of prose and list learning. Journal of Experimental Psychology: General, 114 (2), 213–238.

Rubinstein, D. Y., Weidemann, C. T., Sperling, M. R., & Kahana, M. J. (2023, June). Direct brain recordings suggest a causal subsequent-memory effect. Cerebral Cortex, 33 (11), 6891–6901.

Rudoler, J. H., Herweg, N. A., & Kahana, M. J. (2023, January). Hippocampal theta and episodic memory. Journal of Neuroscience, 43 (4), 613–620. doi: 10.1523/JNEUROSCI.1045-22.2022

Sanquist, T. F., Rohrbaugh, J. W., Syndulko, K., & Lindsley, D. B. (1980). Electrocortical signs of levels of processing: Perceptual analysis and recognition memory. Psychophysiology, 17 (6), 568–576. doi: 10.1111/j.1469-8986.1980.tb02299.x

Satterthwaite, F. E. (1946, December). An approximate distribution of estimates of variance components. Biometrics Bulletin, 2 (6), 110. Retrieved from 10.2307/3002019 doi: 10.2307/3002019

Sederberg, P. B., Kahana, M. J., Howard, M. W., Donner, E. J., & Madsen, J. R. (2003). Theta and gamma oscillations during encoding predict subsequent recall. Journal of Neuroscience, 23 (34), 10809–10814. doi: 10.1523/JNEUROSCI.23-34-10809.2003

Sun, B., Feng, J., & Saenko, K. (2015, December). Return of Frustratingly Easy Domain Adaptation. arXiv.

Wang, C., Subramaniam, V., Yaari, A. U., Kreiman, G., Katz, B., Cases, I., & Barbu, A. (2023). Brainbert: Self-supervised representation learning for intracranial recordings. In Iclr.

Weidemann, C. T., & Kahana, M. J. (2021). Neural measures of subsequent memory reflect endogenous variability in cognitive function. *Journal of Experimental Psychology: Learning*, Memory, and Cognition, 47 (4), 641–651. doi: 10.1037/xlm0000966

Weidemann, C. T., Kragel, J. E., Lega, B. C., Worrell, G. A., Sperling, M. R., Sharan, A. D., … Kahana, M. J. (2019). Neural activity reveals interactions between episodic and semantic memory systems during retrieval. Journal of Experimental Psychology: General, 148 (1), 1–12. doi: 10.1037/xge0000480

Xie, W., Bainbridge, W. A., Inati, S. K., Baker, C. I., & Zaghloul, K. A. (2020). Memorability of words in arbitrary verbal associations modulates memory retrieval in the anterior temporal lobe. Nature Human Behaviour, 4, 937–948.

